# The genetic basis of natural variation in *C. elegans* telomere length

**DOI:** 10.1101/051276

**Authors:** D.C. Cook, S. Zdraljevic, R.E. Tanny, B. Seo, D.D. Riccardi, L.M. Noble, M.V. Rockman, M.J. Alkema, C. Braendle, J.E. Kammenga, J. Wang, L. Kruglyak, M.A. Félix, J. Lee, E.C. Andersen

## Abstract

Telomeres are involved in the maintenance of chromosomes and the prevention of genome instability. Despite this central importance, significant variation in telomere length has been observed in a variety of organisms. The genetic determinants of telomere-length variation and their effects on organismal fitness are largely unexplored. Here, we describe natural variation in telomere length across the *Caenorhabditis elegans* species. We identify a large-effect variant that contributes to differences in telomere length. The variant alters the conserved oligosaccharide/oligonucleotide-binding fold of POT-2, a homolog of a human telomere-capping shelterin complex subunit. Mutations within this domain likely reduce the ability of POT-2 to bind telomeric DNA, thereby increasing telomere length. We find that telomere-length variation does not correlate with offspring production or longevity in *C. elegans* wild isolates, suggesting that naturally long telomeres play a limited role in modifying fitness phenotypes in *C. elegans*.

## Introduction

Genome-wide association (GWA) studies, in which phenotypic differences are correlated with genome-wide variation in populations, offer a powerful approach to understand the genetic basis of complex traits (McCarthy *et al.* 2008). GWA requires accurate and quantitative measurement of traits for a large number of individuals. Even in organisms that are studied easily in the laboratory, the measurement of quantitative traits is difficult and expensive. By contrast, the rapid decrease in sequencing costs has made the collection of genome-wide variation accessible. From *Drosophila* (Mackay *et al.* 2012; Lack *et al.* 2015) to *Arabidopsis* (Weigel and Mott 2009) to humans (Project *et al.* 2012), the whole genomes from large populations of individuals can be analyzed to identify natural variation that is correlated with quantitative traits. Because the genome itself can vary across populations, whole-genome sequence data sets can be mined for traits without measuring the physical organism. Specifically, large numbers of sequence reads generated from individuals in a species can be analyzed to determine attributes of genomes, including mitochondrial or ribosomal DNA copy numbers. Another such trait is the length of the highly repetitive structures at the ends of linear chromosomes called telomeres (Blackburn 1991).

Telomeres are nucleoprotein complexes that serve as protective capping structures that prevent chromosomal degradation and fusion (O’Sullivan and Karlseder 2010). The DNA component of telomeres in most organisms consists of long stretches of nucleotide repeats that terminate in a single-stranded 3’ overhang (McEachern *et al.* 2000). The addition of telomeric repeats is necessary because DNA polymerase is unable to completely replicate the lagging strand (Watson 1972; Levy *et al.* 1992). The length of telomeres can differ among cell populations (Samassekou *et al.* 2010), from organism to organism (Fulcher *et al.* 2014), and within proliferating cellular lineages (Frenck Jr. *et al.* 1998). Two antagonistic pathways regulate telomere length. In the first pathway, the reverse transcriptase telomerase adds *de novo* telomeric repeats to the 3’ ends of chromosomes. In the second, telomere lengthening is inhibited by the shelterin complex. Shelterin forms a protective cap at telomere ends, presumably through the formation of lariat structures known as t-loops (Griffith *et al.* 1999). The t-loops are hypothesized to inhibit telomerase activity by preventing access to the 3’ tail. Additionally, because uncapped telomeres resemble double-stranded DNA breaks, shelterin association with telomeric DNA represses endogenous DNA damage repair pathways, preventing chromosomal fusion events and preserving genome integrity (De Lange 2010).

Variation in telomere length has important biological implications. In cells lacking telomerase, chromosome ends become shorter with every cell division, which eventually triggers cell-cycle arrest (Harley *et al.* 1992). In this way, telomere length sets the replicative potential of cells and acts as an important tumor-suppressor mechanism (Harley *et al.* 1992; Deng *et al.* 2008). In populations of non-clonal human leukocytes, telomere lengths have been shown to be highly heritable (Broer *et al.* 2013). Quantitative trait loci (QTL) identified from human genome-wide association (GWA) studies of telomere length implicate telomere-associated genes, including telomerase (*TERT*), its RNA template (*TERC*), and *OBFC1* (Levy *et al.* 2010; Jones *et al.* 2012; Codd *et al.* 2013). QTL underlying variation in telomere length have been identified in *Arabidopsis thaliana*, *Saccharomyces paradoxus*, and *Saccharomyces cerevisiae* using both linkage and association approaches (Gatbonton *et al.* 2006; Liti *et al.* 2009; Kwan *et al.* 2011; Fulcher *et al.* 2014). In *S. paradoxus*, natural variation in telomere lengths is mediated by differences in telomerase complex components. In *S. cerevisiae*, natural telomere lengthening is caused by a loss of an amino acid permease gene. Thus far, no studies in multicellular animals or plants have been able to identify specific genes responsible for population telomere-length differences. Recent advances in wild strain genotypes and sequences (Andersen *et al.* 2012) in *Caenorhabditis elegans* make it a powerful model to address natural variation in telomere length and its fitness consequences.

Like in humans, telomerase and shelterin activities regulate *C. elegans* telomere length (Malik *et al.* 2000; Cheung *et al.* 2006; Meier *et al.* 2006; Shtessel *et al.* 2013). The TRT-1-containing telomerase complex is hypothesized to add TTAGGC repeats to the ends of chromosomes and prevents chromosome shortening (Meier *et al.* 2006), and the shelterin complex regulates access of the telomerase complex to chromosome ends (Raices *et al.* 2008; Cheng *et al.* 2012; Shtessel *et al.* 2013). The length of telomeres in the laboratory strain N2 is variable and ranges between 2-9 kb (Wicky *et al.* 1996; Raices *et al.* 2005). The telomere lengths in wild isolates of *C. elegans* are largely unexplored. Previous studies examined variation in telomere length using a small number of wild strains (Cheung *et al.* 2004; Raices *et al.* 2005). However, several of the supposed wild strains have since been determined to be mislabeled versions of the laboratory strain N2 (McGrath *et al.* 2009). Thus, it is not known if and how telomere lengths vary among *C. elegans* natural strains. Additionally, the fitness consequences of telomere length variation have not been defined.

Here, we collected a new set of whole-genome sequences from 208 wild *C. elegans* strains and used these strains to investigate natural variation in telomere length across the species. Computational estimates of telomere lengths were confirmed using molecular measurements, indicating that the computational technique can be applied across this large number of wild strains. Using association mapping, we found that variation in the gene *pot-2* is correlated with differences in *C. elegans* telomere length. Natural variation in *pot-2* affects gene function and causes longer than average telomeres in some wild strains. Additionally, we examined whether population differences in telomere length connect to differences fitness traits, including brood size and longevity. Our results indicate that variation in *pot-2* does not correspond with variation in fitness as measured in the laboratory and does not show strong signatures of selection in nature. These data suggest that telomere length beyond a basal threshold is of limited consequence to *C. elegans*. Our results underscore how traits obtained from sequence data can be utilized to understand the dynamic nature of genomes within populations.

## Results

### Whole-genome sequencing of a large number of wild *C. elegans* strains identifies new isotypes and highly diverged strains

Previous genome-wide analyses of *C. elegans* population diversity used single-nucleotide variants (SNVs) ascertained from only two strains (Rockman and Kruglyak 2009), reduced representation sequencing that only studied a fraction of the genome (Andersen *et al.* 2012), or were limited to a small set of wild strains (Thompson *et al.* 2013). To address these limitations, we sequenced the whole genomes of a collection of 208 wild strains (Supplementary File 1). Because *C. elegans* reproduction occurs primarily through the self fertilization of hermaphrodites, highly related individuals proliferate in close proximity to one another (Barrière and Félix 2005; Félix and Braendle 2010). As a result, strains isolated from similar locations in nature are frequently identical and share genome-wide haplotypes or isotypes. Sequencing data generated from strains belonging to the same isotype can be combined to increase depth of coverage and to improve downstream analyses. To identify which strains shared the same genome-wide haplotypes, we compared all of the variation identified in each of the 208 strains to each other in pairwise comparisons. The 208 strains reduce to 152 unique genome-wide haplotypes or isotypes (Supplementary File 1). The combination of sequence data from all strains that make up an isotype led to a 70-fold median depth of coverage (Supplementary Figure 1), enabling the discovery of single nucleotide variants (SNVs) and other genomic features. The number of SNVs in comparisons of each isotype to the reference strain N2 ranged from strains highly similar to N2 to 402,436 SNVs (Supplementary Figure 2), and the density of SNVs across the genome matched previous distributions with more variants on chromosome arms than centers (Supplementary Figure 3). A clustering analysis of these 152 isotypes recapitulated the general relationships previously identified among a set of 97 wild isotypes (Andersen *et al.* 2012) (Supplementary Figure 4) with the addition of 55 new isotypes. Past studies identified one highly diverged strain isolated from San Francisco, QX1211, which had divergence almost three times the level of other wild *C. elegans* strains (Andersen *et al.* 2012). Among the 55 new isotypes, one additional strain, ECA36 from New Zealand, is equally diverged, suggesting that wider sampling will recover additional diversity for this species. Altogether, our considerably expanded collection of whole-genome sequence data serves as a powerful tool to interrogate how natural variation gives rise to differences among individuals in a natural population.

### *C. elegans* wild strains differ in telomere lengths

Our collection of high-depth whole-genome sequence data samples a large number of strains in the *C. elegans* species. The recent development of TelSeq, a program designed to estimate telomere length using short-read sequence data (Ding *et al.* 2014), allowed us to examine natural variation in telomere lengths computationally across wild *C. elegans* strains. We detected considerable natural variation in the total length of telomeric DNA in a strain (Figure 1), ranging from 4.12 kb to a maximum of 83.7 kb and a median telomere length of 12.25 kb (Supplementary File 2). The Telseq telomere length estimate for N2 was 16.97 kb, which is higher than previous estimates (Wicky *et al.* 1996). The distribution of telomere lengths in the *C. elegans* population approximated a normal distribution with a right tail containing strains with longer than average telomeres. We found that our computational estimates of telomere length from Illumina sequence data were significantly influenced by library preparation, possibly driven by the method of DNA fragmentation (Supplementary Figure 5). However, we were able to control for these differences using a linear model. We also observed a weak correlation between depth of coverage and TelSeq length estimates, but adjustments for library preparation eliminated this relationship (Supplementary Figure 6).

**Fig 1.**
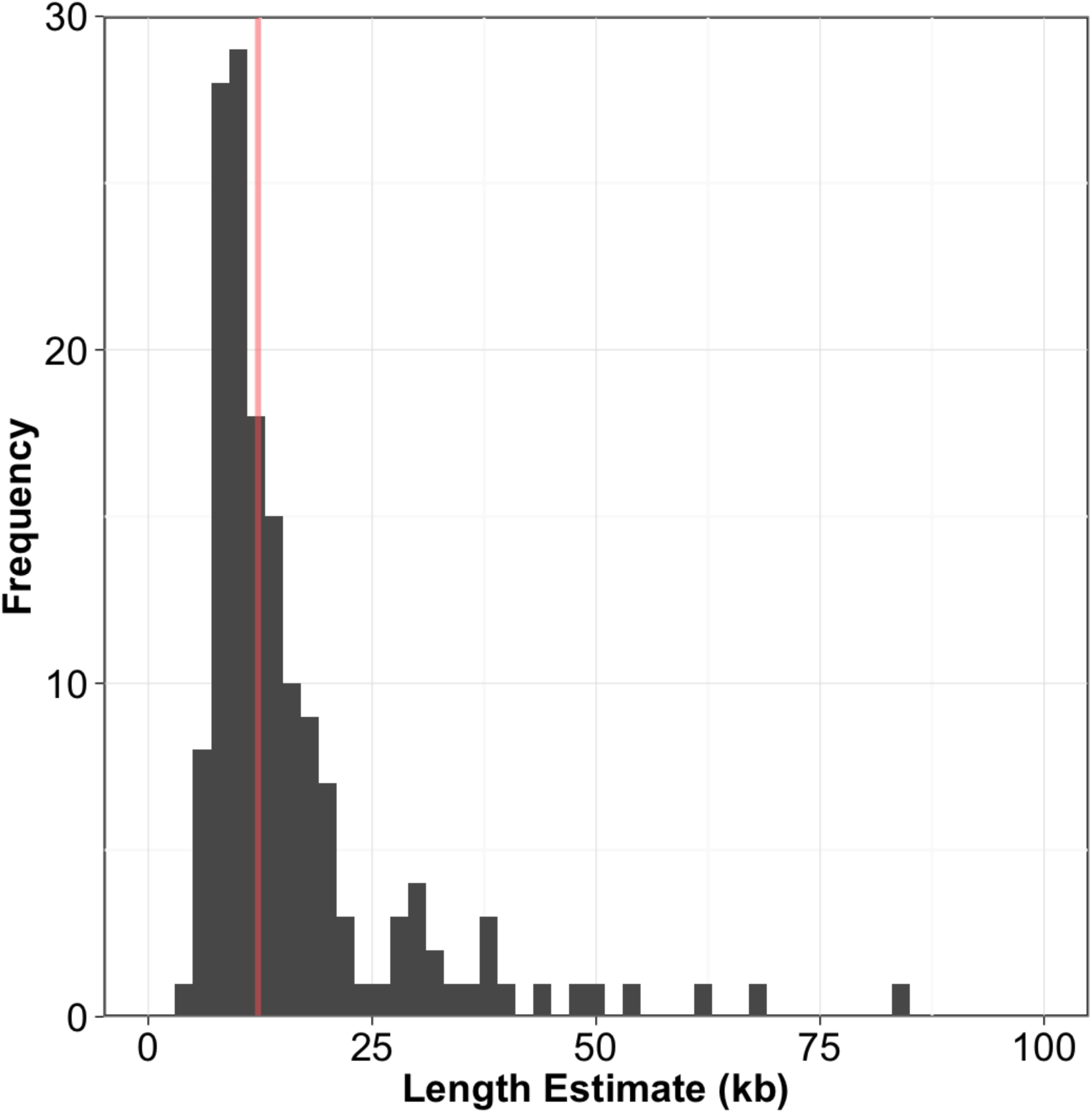
Distribution of telomere-length estimates. A histogram of telomere-length estimates weighted by the number of reads sequenced per run is shown. Bin width is 2. The red line represents the median telomere-length estimate of 12.2 kb.

TelSeq length estimates have been shown to give similar results as molecular methods to measure human telomere length (Ding *et al.* 2014). As of now, no studies have used TelSeq to examine *C. elegans* telomeres, so we investigated how well TelSeq estimates correlated with molecular methods, including Terminal Restriction Fragment (TRF) Southern blot analyses, quantitative PCR (qPCR) of telomere hexamer sequences, and fluorescence *in situ* hybridization (FISH) analyses. Using twenty strains, we found that the results from these molecular assays correlated well (*rho* = 0.445 TRF, 0.815 FISH, 0.699 qPCR, Spearman’s rank correlation) with computational estimates of telomere lengths (Figure 2, Supplementary File 3). These molecular results validated our computational estimates of telomere lengths and indicate that we can use TelSeq estimates to investigate the genetic causes underlying telomere variation.

**Fig 2.**
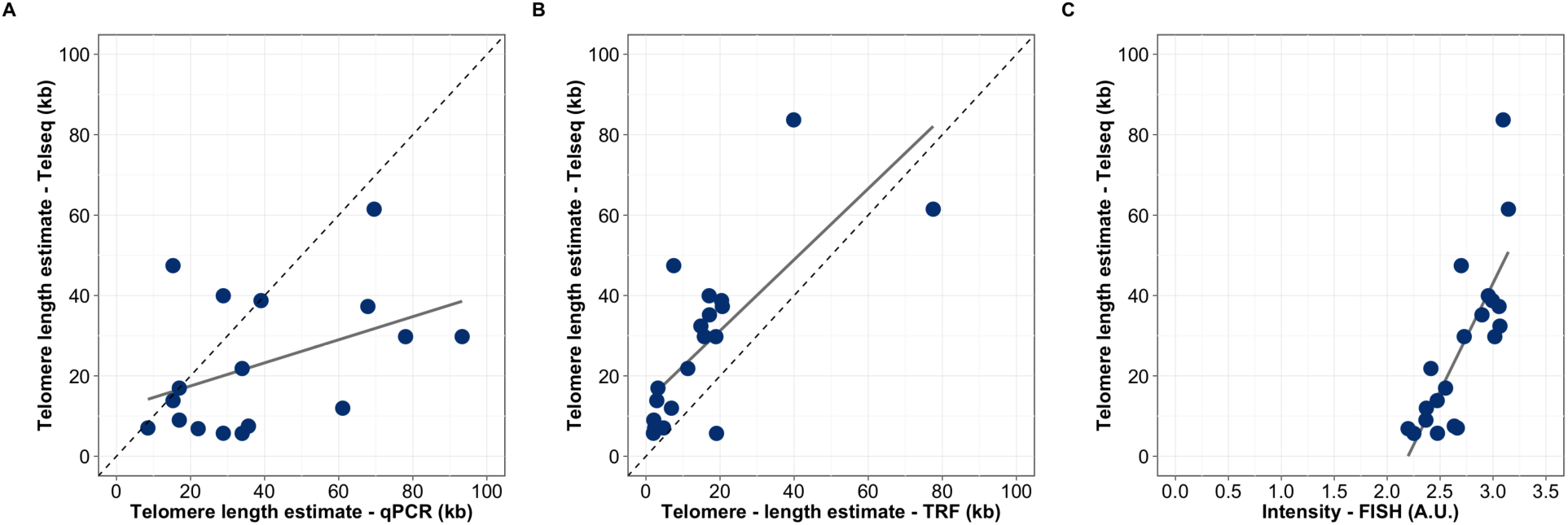
Telomere-length estimates correlate with alternative molecular measurement methods. Scatterplot of Telseq Telomere-length estimates (y-axis) plotted against alternative methods of telomere length measurement on the x-axis. Alternative methods plotted on the x-axis and their associated Spearman’s rank correlation are (A) qPCR measurements normalized by N2 qPCR value and scaled relative to the TelSeq N2 telomere length estimate (*rho* = 0.445, p = 0.049), (B) TRF (*rho* = 0.699, p = 8.5e^−4^) and (C) FISH (*rho* = 0.815, p = 1.03e^−5^). Grey lines represent the regression lines between Telseq and each method. Dashed diagonal lines represent identity lines.

### Species-wide telomere length differences correlate with genetic variation on chromosome II

To identify the genes that cause differences in telomere length across the *C. elegans* population, we used a GWA mapping approach as performed previously (Andersen *et al.* 2012) but taking advantage of the larger collection of wild strains. We treated our computational estimates of telomere length as a quantitative trait and identified one significant quantitative trait loci (QTL) on the right arm of chromosome II (Figure 3, Supplementary File 4). To identify the variant gene(s) that underlie this QTL, we investigated the SNVs within a large genomic region (12.9 to 15.3 Mb) surrounding the most significant marker on chromosome II. This region contains 557 protein-coding genes (Supplementary File 5), but only 332 of these genes contained variants that are predicted to alter the amino-acid sequences among the 152 strains. We examined genes with predicted protein coding variants that could alter telomere length by correlating their alleles with the telomere-length phenotype. Thirty-four genes possessed variation that was most highly correlated with telomere length (*rho* ≥ 0.4, Supplementary File 5). The chromosome II QTL explains 28.4% of the phenotypic variation in telomere length. Three additional suggestive QTL on chromosomes I, II, and III were detected close to but below the significance threshold. Taken together, the four QTL explain 56.7% of the phenotypic variation in telomere length.

**Fig 3.**
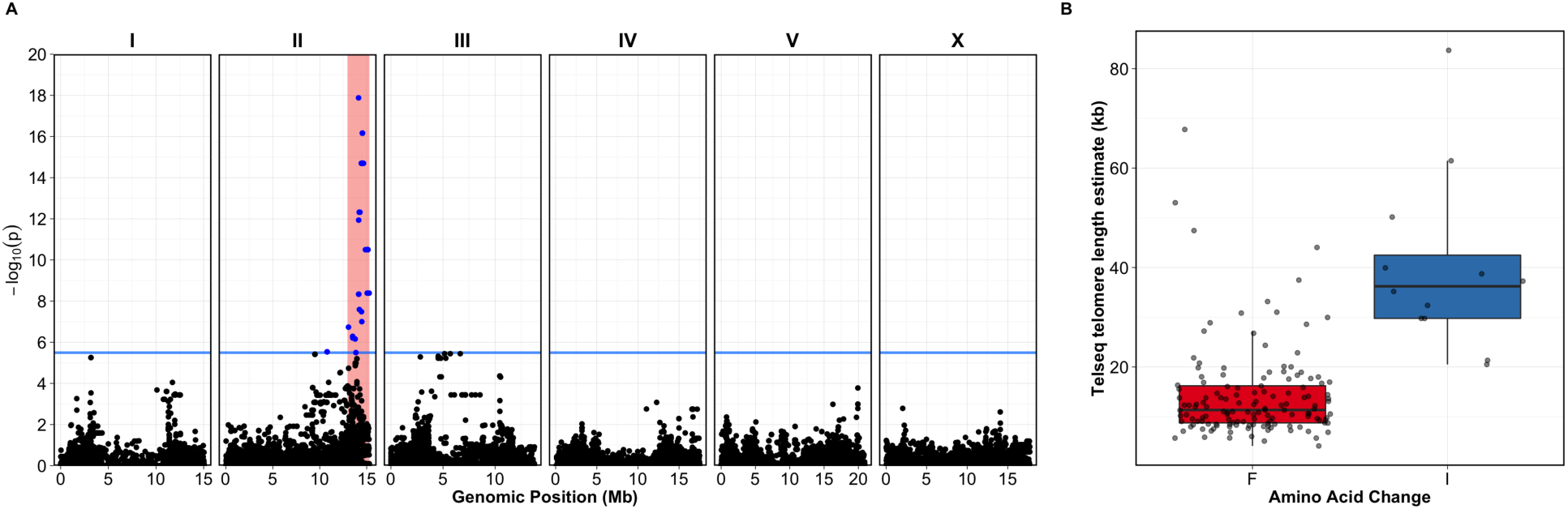
Genome-wide association of telomere length. (A) Genome-wide association of telomere length residuals (conditioned on DNA library) is visualized using a Manhattan plot. Genomic coordinates are plotted on the x-axis against the negative of the log-transformed *p*-value of a test of association on the y-axis. The blue bar indicates the Bonferroni-corrected significance threshold (*α* = 0.05). Blue points represent SNVs above the significance threshold whereas black points represent SNVs below the significance threshold. Light-red regions represent the confidence intervals surrounding significantly associated peaks. (B) Shown is the split between TelSeq estimated telomere lengths (y-axis) by genotype of *pot-2* at the presumptive causative allele as boxplots (x-axis). The variant at position 14,524,396 on chromosome II results in a putative F68I coding change. Horizontal lines within each box represent the median, and the box represents the interquartile range (IQR) from the 25th-75th percentile. Whiskers extend to 1.5x the IQR above and below the box. The plotted points represent estimates beyond 1.5x the IQR.

### Variation in *pot-2* underlies differences in telomere length

One of the 34 genes in the chromosome II large-effect QTL is *pot-2* (Protection Of Telomeres 2), a gene which was previously implicated in regulation of telomere length (Raices *et al.* 2008; Cheng *et al.* 2012; Shtessel *et al.* 2013). Given the large number of genes present within our confidence interval and challenges associated with examining telomere length using traditional genetic approaches, we sought alternative methods to confirm that variation in *pot-2* could cause long telomeres. A quantitative complementation test could be used to confirm that wild strains have the same functional effect as a *pot-2* deletion. However, differences in telomere length caused by mutations in genes that encode telomere-associated proteins often do not have observable telomere defects for a number of generations (Vulliamy *et al.* 2004; Armanios *et al.* 2005; Marrone *et al.* 2005). It is technically not feasible to keep the genome heterozygous during long-term propagation. Fortunately, the ability to computationally estimate telomere length allowed us to further validate our approach using data from the Million Mutation Project (MMP) (Thompson *et al.* 2013) and examine whether the equivalent of a mutant screen for telomere length would provide insight into our result examining wild isolate genomes.

The MMP generated over two thousand mutagenized strains using the laboratory N2 background. After each strain was passaged by self-mating of hermaphrodites for ten generations, the strains were whole-genome sequenced to identify and to predict the effects of induced mutations. The MMP data set can be used to identify correlations of phenotype and mutant genes in the laboratory strain background. We obtained whole-genome sequence data from 1,936 mutagenized N2 strains, each of which has a unique collection of mutations. Importantly, ten generations of self-propagation of these mutagenized strains prior to sequencing likely allowed telomere lengths to stabilize in response to mutations in genes that regulate telomere length, enabling us to observe differences. TelSeq returned telomere length estimates for this population, which had a right long-tailed distribution (Figure 4A). We classified 39 of 1936 strains within the population as long-telomere strains with telomere lengths greater than 6.41 kb (98th percentile). Reasoning that certain mutant genes would be overrepresented in these 39 strains compared to the others, we performed a hypergeometric test to identify if enrichment for particular genes in long-telomere strains existed. After adjusting for multiple statistical tests, we identified *pot-2* as highly enriched for mutations in six of the 39 long-telomere strains (p = 2.69e^−11^, Bonferroni corrected, Figure 4B). No other genes within any of the QTL intervals or any other part of the genome were enriched for mutations among long-telomere strains. This approach was different from association mapping and identified the same locus regulating telomere length. Additionally, we computationally examined telomere length from whole-genome sequencing of a *pot-2* knockout strain. This strain possesses a large deletion that spans the first and second exons of *pot-2* likely rendering it nonfunctional. We propagated this mutant strain for ten generations prior to whole-genome sequencing and TelSeq analysis. The telomere length of *pot-2(tm1400)* mutants was calculated to be 30.62 kb. Given these data, we have three independent tests that indicate that variation in *pot-2* likely underlies natural differences in telomere lengths across the *C. elegans* species.

**Fig 4.**
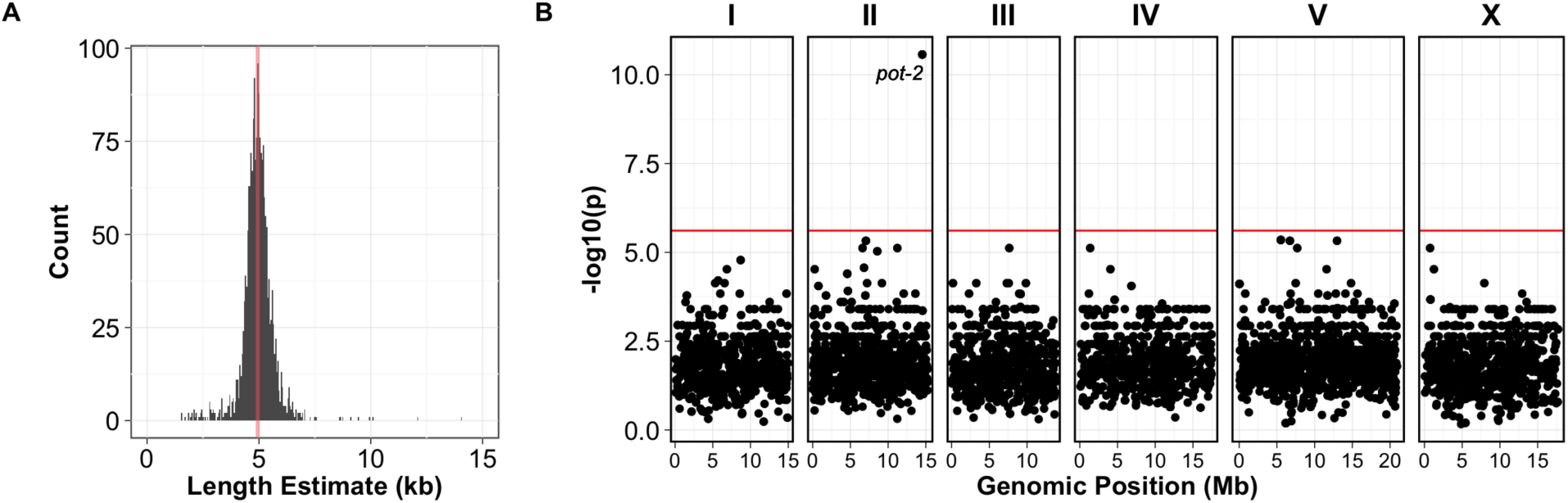
Mutations in *pot-2* are more often found in strains with long telomeres than in strains with short telomeres. (A) A histogram of telomere-length estimates among the 1,936 mutagenized strains from the Million Mutation Project. Median telomere length is 4.94 kb. (B) Plot of significance from a hypergeometric test for every C. elegans protein-coding gene. The red line represents the Bonferroni (*α* = 0.05) threshold set using the number of protein coding genes (20,447). Each point represents a gene plotted at its genomic position on the x-axis, and the log transformed *p*-value testing for enrichment of mutations in long-telomere strains.

Our results are consistent with the established role of *pot-2* as an inhibitor of telomere lengthening (Shtessel *et al.* 2013). However, no connection of *pot-2* to natural variation in telomere lengths has been described previously. POT-2 contains an OB-fold thought to interact with telomeric DNA. OB-folds are involved in nucleic acid recognition (Flynn and Zou 2010), and the OB-fold of the human POT-2 homolog (hPOT1) binds telomeric DNA (Lei *et al.* 2004). We investigated the variant sites altered in the *C. elegans* species along with the mutations found in the MMP mutagenized strains (Figure 5). We found that the natural variation in *pot-2* resulted in a putative phenylalanine-to-isoleucine (F68I) change in the OB-fold domain of 12 strains. Strains with the POT-2(68I) allele have long telomeres on average, whereas strains with POT-2(68F) allele have normal length telomeres on average. Synonymous variants or variation outside of the OB-fold domain were rarely found in strains with long telomeres. Because loss of *pot-2* is known to cause long telomeres (Raices *et al.* 2008; Cheng *et al.* 2012; Shtessel *et al.* 2013), the F68I variant likely reduces or eliminates the function of *pot-2*. Additionally, six out of the 39 long-telomere MMP strains had mutations in *pot-2*, including five strains that had mutations within or directly adjacent to the OB-fold and an additional strain with a nonsense mutation outside the OB-fold domain that likely destabilizes the transcript. These data support the hypothesis that *pot-2* is the causal gene underlying variation in telomere lengths across the *C. elegans* species.

**Fig 5.**
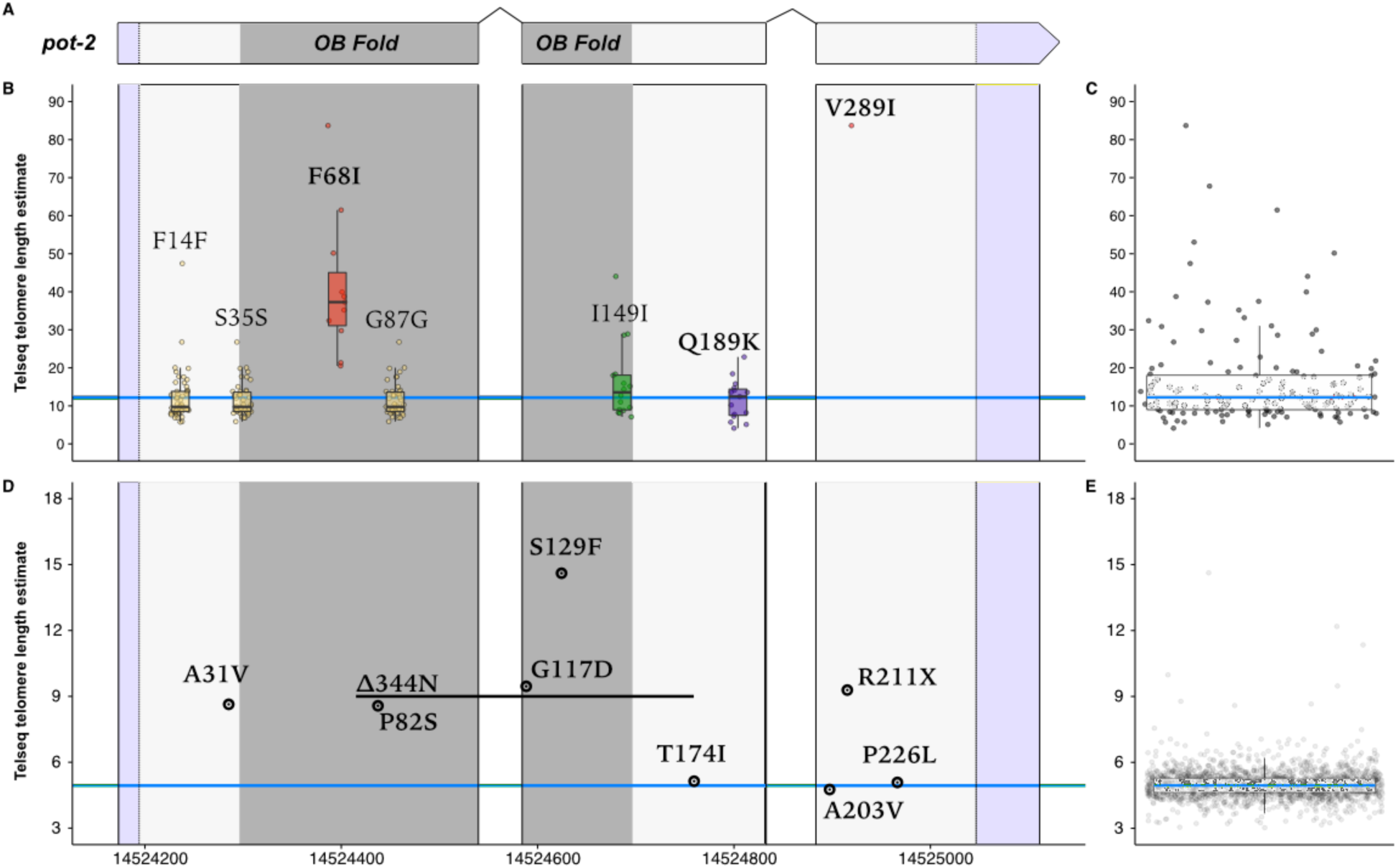
Variation within *pot-2* in wild isolate and Million Mutation Project strains. Natural variation and induced mutations that alter codons across *pot-2* are shown along with the telomere-length estimates for all strains. In panel (A), a schematic illustrating the *pot-2* genomic region is shown. The dark gray region represents the part of the genome encoding the OB-fold domain. Purple regions represent untranslated regions. (B) Strains that harbor the alternative (non-reference) allele are plotted by telomere length on the y-axis and genomic position on the x-axis. Both synonymous and nonsynonymous variants are labeled. Variants resulting in a nonsynonymous coding change are bolded. The blue line indicates the median telomere length value for wild isolates. The color of boxplots and markers indicates variants from the same haplotypes. (C) Boxplot of natural isolate distribution of telomere lengths. Blue lines within the center of each box represent the median while the box represents the interquartile range (IQR) from the 25th – 75th percentile. Whiskers extend to 1.5x the IQR above and below the box. All plotted points represent estimates beyond 1.5x the IQR. (D) Telomere length is plotted on the y-axis as in (B), but strains do not share mutations because strains harbor unique collections of induced alleles. The blue line indicates median telomere length for the MMP population. (E) Boxplot of the distribution of telomere lengths in the MMP is shown. Boxplot follows same conventions as in (C). N2 telomere length in our wild isolate population was estimated to be 16.9 kb whereas median telomere length in MMP was 4.94 kb. Differences are likely caused by library preparation method and/or sequencing method.

### Natural variants in *pot-2* do not have detectable fitness consequences

We connected genetic variation in the gene *pot-2* with telomere-length differences across *C. elegans* wild strains. Specifically, an F68I variant in the putative telomere-binding OB-fold domain might cause reduction of function and long telomeres. A variety of studies have observed a relationship between telomere length and organismal fitness, including longevity or cellular senescence (Harley *et al.* 1992; Heidinger *et al.* 2012; Soerensen *et al.* 2012). Our results with natural variation in telomere lengths provided a unique opportunity to connect differences in the length of telomeres with effects on organismal fitness. We measured offspring production for our collection of 152 wild strains and found no correlation with telomere length (*rho*=0.062, Figure 6A). Long telomeres allow for increased replicative potential of cells (Harley *et al.* 1992), but it is unclear how the replicative potential of individual cells contributes to organismal longevity phenotypes (Hornsby 2007). We chose nine strains covering the range of telomere-length differences and found no correlation with longevity (*rho*=-0.008, Figure 6B, Supplementary Figure 7). Taken together, these results suggest that the long telomeres found in some wild *C. elegans* strains do not have significant fitness consequences in these laboratory-based experiments.

**Fig 6.**
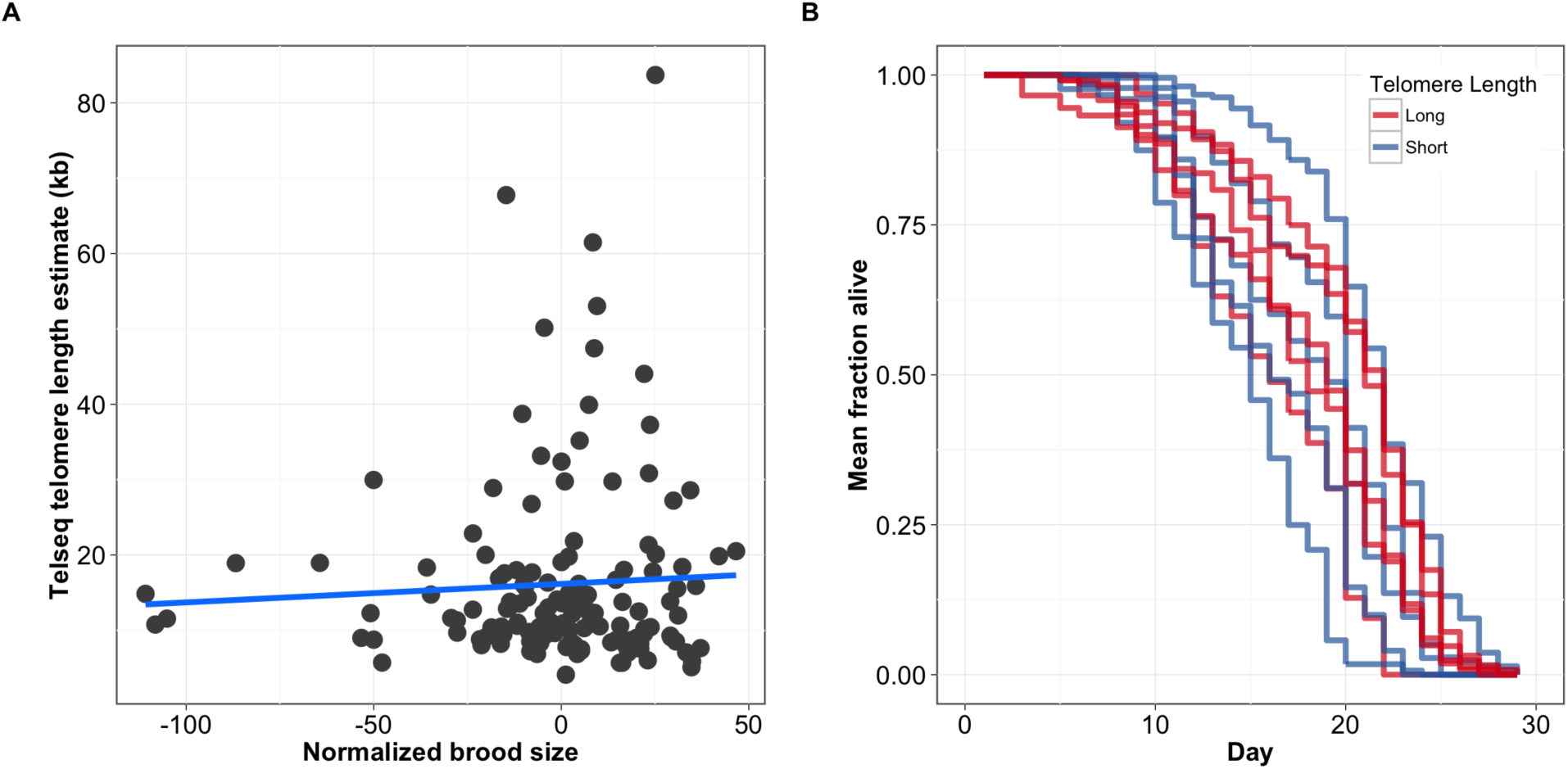
Fitness traits are not associated with telomere length. (A) Normalized brood sizes (x-axis) of 152 wild isolates are plotted against the telomere-length estimates from those same strains (y-axis). The blue line indicates a linear fit of the data. However, the correlation is not significant (*rho* = -0.062, p = 0.463). (B) Survival curves of nine wild isolates with long and short telomeres. Lines represent aggregate survival curves of three replicates. Survival among long and short telomere length strains is not significantly different (p = 0.358; Mantel-Cox analysis).

Because we did not observe a strong effect on organismal fitness, we investigated the population genetics of *pot-2* to test whether that locus had any signature of selection. Examination of Tajima’s D at the *pot-2* locus yielded no conspicuous signature, though the characteristic high linkage disequilibrium of *C. elegans* makes gene-focused tests challenging in this species (Supplementary Figure 8). Furthermore, the haplotypes that contain this variant are rare (Supplementary Figure 9) and not geographically restricted (Supplementary Figure 10) Like the measurements of organismal fitness and lack of correlation with telomere-length differences, the population genetic measures for selection indicate that the observed variation in *pot-2* is not under strong selective pressure. Together, these results suggest that natural variation in telomere length plays a limited role in modifying whole-organism phenotypes in *C. elegans*.

## Discussion

In this study, we report the identification of a QTL on the right arm of chromosome II containing a variant within the gene *pot-2* that contributes to differences in telomere length of *C. elegans* wild isolates. Several lines of evidence support the F68I allele of *pot-2* as the variant modulating telomere lengths. First, others have shown previously that loss of *pot-2* results in progressive telomere lengthening in the laboratory strain background (Raices *et al.* 2008; Shtessel *et al.* 2013). Additionally, the F68I variant is the only SNV in *pot-2* that correlates with long telomeres. This variant falls within the OB-fold of POT-2, and our examination of strain telomere lengths within the MMP shows enrichment of mutations from long-telomere strains found within the OB-fold domain. Importantly, OB-folds are known to interact with single-stranded nucleic acids, and TelSeq telomere-length estimates of wild isolates and randomly mutagenized laboratory strains show that mutation or variation of the OB-fold domain reduces function and causes long telomeres, as we also observed in the *pot-2(tm1400)* deletion strain. Moreover, this amino acid change could plausibly alter the function of the OB-fold within POT-2. Nucleic acid recognition of OB-folds occurs through a variety of molecular interactions, including aromatic stacking (Gatzeva-topalova *et al.* 2011). A change from phenylalanine to isoleucine would eliminate a potential aromatic stacking interaction and presumably reduce the binding affinity and function of POT-2. Somatic mutations in individuals with chronic lymphocytic leukemia were found to be concentrated within the OB-folds of hPOT1 (Ramsay *et al.* 2013).

We wondered why additional genes involved in the regulation of telomeres were not identified from our study of telomere lengths across wild isolates and mutagenized laboratory strains. Homologs for both telomerase and shelterin complex components are found in *C. elegans* (Stein *et al.* 2001). We identified natural variation in *trt-1* but only in the highly diverged strains ECA36 and QX1211. These rare alleles are removed from the GWA mapping, because we require allele frequencies to be greater than 5%. Laboratory mutants in *trt-1* have short telomeres (Cheung *et al.* 2006; Meier *et al.* 2006), but we do not see enrichment of *trt-1* mutations in the MMP collection for short or long telomeres. *C. elegans* contains orthogolous genes for two of the six shelterin complex members, hPOT1 and RAP1 (Harris *et al.* 2009). Four *C. elegans* genes with homology to hPOT1 have been identified (*mrt-1*, *pot-1*, *pot-2*, and *pot-3*) (Raices *et al.* 2008; Meier *et al.* 2009), and *C. elegans rap-1* is homologous to human RAP1 (Raices *et al.* 2008; Meier *et al.* 2009). The genes *rap-1* and *pot-3* had no variants or only rare variants, respectively. All of the other homologous genes contained variants in 5% or more of the wild isolates. None of these genes mapped by GWA besides *pot-2*, and none of the mutations in these genes were enriched in short-or long-telomere strains from the MMP collection. Perhaps shorter telomere strains are less fit and do not survive well in the wild or during the growth of mutant MMP strains. These results suggest that long telomeres are likely of limited consequence compared to short telomeres in natural settings. Additionally, because TelSeq provides an average estimate of telomere length, it is possible for the variance of telomere lengths to increase without affecting average length estimates. For this reason, we might not detect a QTL at *pot-1*, which has been previously reported to result in longer but more heterogeneous telomeres (Raices *et al.* 2008).

Our observation that considerable telomere-length variation in the wild isolate population exists allowed us to directly test whether variation in telomere length contributes to organismal fitness. We did not see any correlation between telomere length and offspring production, suggesting that fitness in wild strains is not related to telomere length. In contrast to findings in human studies, we did not identify a relationship between telomere length and longevity. Our results confirm past findings that telomere length is not associated with longevity in a small number of *C. elegans* wild isolates or laboratory mutants (Raices *et al.* 2005). Although the effects of telomere length on longevity have been observed in a well controlled study of the gene *hrp-1* on isogenic populations in the laboratory (Joeng *et al.* 2004), this study differs from our results in wild isolates. The background effects of wild isolate variation along with telomere-length variation could obfuscate a direct connection to longevity. Even though we did not identify a correlation between telomere length and either longevity or offspring production under laboratory conditions, our study suggests a limited role for telomeres in post-mitotic cells. Furthermore, the population genetic results do not strongly support evidence of selection on *pot-2* variants.

In summary, this study demonstrates that a variant in *pot-2* likely contributes to phenotypic differences in telomere length among wild isolates of *C. elegans*. The absence of evidence for selection at the *pot-2* locus and the lack of strong effects on organismal fitness traits suggest that differences in telomere length do not substantially affect individuals at least under laboratory growth conditions. Additionally, our study demonstrates the ability to extract and to utilize phenotypic information from sequence data. A number of approaches can be employed to examine other dynamic components of the genome, including mitochondrial and ribosomal DNA copy numbers, the mutational spectrum, or codon biases. These traits present a unique opportunity to identify how genomes differ among individuals and the genetic variants underlying those differences.

## Materials and Methods

### Strains

*C. elegans* strains were cultured using bacterial strain OP50 on a modified nematode growth medium (NGMA, 1% agar, 0.7% agarose) to prevent burrowing of wild isolates (Andersen *et al.* 2014). Strain information is listed in Supplementary File 1. The following strains were scored for the molecular telomere assays described below: AB4 (CB4858 isotype), CB4856, CX11285, CX11292, DL238, ECA248, ED3012, EG4349, JT11398, JU311, JU1400, JU2007, KR314, N2, NIC2, NIC3, NIC207, PB303, and QX1212.

### Library construction and sequence acquisition

DNA was isolated from 100-300 *μ*l of packed animals using the Blood and Tissue DNA isolation kit (Qiagen). The provided protocol was followed with the addition of RNAse (4 *μ*l of 100 mg/ml) following the initial lysis for two minutes at room temperature. DNA concentration was determined using the Qubit dsDNA BR Assay Kit (Invitrogen). Libraries were generated using the Illumina Nextera Sample Prep Kit and indexed using the Nextera Index Kit. Twenty-four uniquely indexed samples were pooled by mixing 100 ng of each sample. The pooled material was size-selected by electrophoresing the DNA on a 2% agarose gel and excising the fragments ranging from 300-500 bp. The sample was purified using the Qiagen MinElute Kit and eluted in 11 *μ*l of buffer EB. The concentration of the purified sample was determined using the Qubit dsDNA HS Assay Kit. Sequencing was performed on the Illumina HiSeq 2500 platform. To increase coverage of some strains, we incorporated data from two separate studies of wild strains (Thompson *et al.* 2013; Noble *et al.* 2015).

### Trimming and demultiplexing

When necessary, demultiplexing and sequence trimming were performed using fastx_barcode_splitter.pl (version 0.0.14) (Gordon and Hannon 2010). Sequences were trimmed using trimmomatic (version 0.32) (Bolger *et al.* 2014). Nextera libraries were trimmed using the following parameters:

> NexteraPE-PE.fa:2:80:10 MINLEN:45

TruSeq libraries were trimmed using:

> TruSeq2-PE.fa:2:80:10 TRAILING:30 SLIDINGWINDOW:4:30 MINLEN:30

The full details of the preparation, source, and library are available in Supplementary File 6.

### Alignment, variant calling, and filtering

FASTQ sequence data has been deposited under NCBI Bioproject accession PRJNA318647. Sequences were aligned to WS245 (http://www.wormbase.org) using BWA (version 0.7.8-r455) (Li and Durbin 2009). Optical/PCR duplicates were marked with PICARD (version 1.111). BAM and CRAM files are available at www.elegansvariation.org/Data. To determine which type of SNV caller would perform best on our dataset and to set appropriate filters, we simulated variation in the N2 background. We used <u>bamsurgeon</u> (github.com/adamewing/bamsurgeon), which modifies base calls to simulate variants at specific positions within aligned reads and then realigns reads to the reference genome using BWA. We simulated 100,000 SNVs in 10 independent simulation sets. Of the 100,000 sites chosen in each simulation set, bamsurgeon successfully inserted an average of 95,172.6 SNVs. Using these 10 simulated variant sets, we tested two different methods of grouping our strains for variant calling: calling strains individually (comparing sequences from a single strain to the reference) or calling strains jointly (comparing all strains in a population to each other). After grouping, bcftools has two different calling methods: a consensus caller (specified using-c), and a more recently developed multiallelic caller (specified using-m) (Li 2011). We performed variant calling using all four combinations of individual/joint calling and the consensus/multiallelic parameters. Because of the hermaphroditic life cycle of *C. elegans*, heterozygosity rates are likely low. Occasionally, heterozygous variants will be called despite skewed read support for reference or alternative alleles. To account for these likely erroneous calls, we performed ‘heterozygous polarization’ using the log-likelihood ratios of reference to alternative genotype calls. When the log-likelihood ratio was less than −2 or greater than 2, heterozygous genotypes were polarized (or switched) to reference genotypes or alternative genotypes, respectively. All other SNVs with likelihood ratios between −2 and 2 were called NA. Following variant calling and heterozygous polarization on resulting calls, we observed increased rates of heterozygous calls using joint methods and decreased true positive rates using our simulation data set (Supplementary File 7, Supplementary File 8). Given *C. elegans* predominantly asexual mode of reproduction, we decided to focus on the individual-based calling method that performed better. Next, we determined the optimal filters to maximize true positive (TP) rates and minimize false positive (FP) and false negative (FN) results using our simulated data (Supplementary File 8). After implementing different combinations of filters, we found that depth (DP), mapping quality (MQ), variant quality (QUAL), and the ratio of high quality alternative base calls (DV) over DP filters worked well (Supplementary Figure 11). Variants with DP <= 10, MQ <= 40, QUAL < 30, and DV/DP < 0.5 were called NA. Using these filters, we called 1.3M SNVs across 152 isotypes. This data set is available at www.andersenlab.org/Research/Data/Cooketal.

### Validation of SNV calling methods

In addition to performing simulations to optimize SNV-calling filters, we compared our whole-genome sequence variant calls with SNVs identified previously in CB4856 (Wicks *et al.* 2001). Out of 4,256 sites we were able to call in regions that were sequenced using Sanger sequencing, we correctly identified 4,223 variants (99.2% of all variants) in CB4856. One true positive was erroneously filtered and two false positives were removed using our filters, and we failed to call the non-reference allele for 30 variants (false negatives).

Additionally, we examined sequence variants with poor parameters in terms of depth, quality, heterozygosity, or modification by our heterozygous polarization filter. We used primer3 (Rozen and Skaletsky 1998) to generate a pair of primers for performing PCR and a single forward primer for Sanger sequencing. We successfully sequenced 73 of 95 sites chosen from several strains. Comparison of variant calls after imputation and filtering yielded 46 true positives (TP) and 14 true negatives (TN). We successfully removed 3/11 false positives (FP) and erroneously filtered two sites that should have been called as non-reference (FN). We also validated the variant responsible for the F68I change in JT11398.

### Identification of clonal sets

Some strains in our original collection were isolated from the same or nearly identical locations. Therefore, we determined if these strains share distinct genome-wide haplotypes or isotypes. To determine strain relatedness, we sequenced and called variants from sequencing runs independently (*e.g.* individual FASTQ pairs) to ensure that strains were properly labeled before and after sequencing. We then combined FASTQ files from sequencing runs for a given strain and examined the concordance among genotypes. Comparison of variants identified among sequencing strains were used to determine whether the strains carried identical haplotypes. We observed that some strains were highly related to each other as compared with the rest of the population. Strains that were greater than 99.93% identical across 1,589,559 sites were classified as isotypes (Supplementary Figure 12). Because LSJ1 and N2 share a genome-wide genotype but exhibit distinct phenotypes (Sterken *et al.* 2015), we treated each strain as a separate isotype. We found the following isotype differences from the previous characterization of a large number of strains. JU360 and JU363 were previously thought to be separate, but highly related, isotypes. We found that, at the genome-wide level and at high depths of coverage, these strains are from the same isotype. Several wild strains isolated before 2000 had different genome-wide haplotypes compared to strains with the same names but stored at the *Caenorhabditis* Genetics Center (CGC). CB4851 from the CGC had a different genome-wide haplotype compared to a strain with the same name from Cambridge, UK. We renamed the CB4851 strain from the CGC as ECA243. By contrast, the version from Cambridge, UK was nearly identical to N2 and not studied further. CB4855 from the CGC has a genome-wide haplotype that matches CB4858, which has a different history and isolation location. Therefore, we cannot guarantee the fidelity of this strain, and it was not studied further. CB4855 from Cambridge, UK is different from the CGC version of CB4855. We gave this strain the name ECA248 to avoid confusion. CB4858 from CGC has a different genome-wide haplotype than CB4858 from Cambridge, UK. Therefore, we renamed CB4858 from Cambridge, UK to ECA252, and it is a separate isotype. The CB4858 from the CGC was renamed ECA251 and is the reference strain from the CB4858 isotype.

### Imputation and variant annotation

Following SNV calling and filtering, some variant sites were filtered. Therefore, we generated an imputed SNV set using beagle (version r1399) (Browning and Browning 2016). This imputed variant set is available at www.andersenlab.org/Research/Data/Cooketal. We used SnpEff (version 4.1g) (Cingolani *et al.* 2012) on this SNV set to predict functional effects.

### Telomere-length estimation

Telomere lengths were estimated using TelSeq (version 0.0.1) (Ding *et al.* 2014) on BAM files derived from wild isolates or MMP strain sequencing. To estimate telomere lengths, TelSeq determines the reads that contain greater than seven telomeric hexamer repeats (TTAGGC for *C. elegans*). Compared to most hexamers, telomeric hexamers can be found tandemly repeated within sequenced reads. TelSeq calculates the relative proportion of reads that appear to be telomerically derived among all sequenced reads and transforms this value into a length estimate using the formula *l* = *t*_*k*_*SC* where *l* is the length estimate and *t*_*k*_ is the abundance of reads with a minimum of *k* telomeric repeats. The value of *s* is the fraction of all reads with a GC composition similar to the telomeric repeat (48-52% for *C. elegans*). The value of *c* is a constant representing the length of 100 bp windows within the reference genome where GC content is equal to the GC content of the telomeric repeat divided by the total number of telomere ends. By default, TelSeq provides length estimates applicable to humans. We found the number of 100 bp windows with a 50% GC content in the WS245 reference genome to be 58,087. We calculated *c* for *C. elegans* as 58,087 *kb* / 12 *telomeres* = 484. This value was used to transform human length estimates to length estimates appropriate to *C. elegans*.

Notably, telomere length estimates are averaged across all chromosomes, as no specific data about any one particular telomere is determined. To assess how well the TTAGGC hexamer distinguishes telomeric reads from non-telomeric reads, we examined the frequencies of non-cyclical permutations of the *C. elegans* hexamer in the N2 laboratory strain using TelSeq (Supplementary Figure 13). We observe that the majority of hexamers examined were not present in more than six copies in a high frequency of reads. By contrast, the reads possessing the telomeric hexamer with seven copies or more were more abundant than any other hexamer. Tandem repeats of the telomeric hexamer are present within the reference genome at the ends of each chromosome and occasionally internally within chromosomes at between 2-71 copies (Supplementary File 9). After running TelSeq on our wild isolates, we removed eight sequencing runs (out of 868 total) that possessed zero reads with 15 or more copies of the telomeric hexamer. These sequencing runs provide additional support for SNV calling but had short read lengths that would underestimate telomere length. We used the weighted average of telomere length estimates for all runs of a given strain based on total reads to calculate telomere-length estimates. Supplementary File 10 details telomere length estimates for every sequencing run.

### Quantitative PCR assays for telomere-length measurements

Telomere lengths were measured by qPCR as described previously with some modifications (Cawthon 2009). Primer sequences were modified from the vertebrate telomere repeat (TTAGGG) to use the *C. elegans* telomere repeat (TTAGGC):

> telG: 5’-ACACTAAGCTTTGGCTTTGGCTTTGGCTTTGGCTTAGTCT-3’
>
> telC: 5’-TGTTAGGTATGCCTATGCCTATGCCTATGCCTAT**GCCTAA**GA-3’

The internal control, *act-1*, was amplified using the following primer pair:

> forward: 5’-GTCGGTATGGGACAGAAGGA-3’
>
> reverse: 5’-GCTTCAGTGAGGAGGACTGG-3’

Two primer pairs were amplified separately (singleplex qPCR). All the samples were run in triplicate. qPCR was performed using iQ SYBR green supermix (BIORAD) with iCycler iQ real-time PCR detection system (BIORAD). After thermal cycling, C_t_ (cycle thresholds) values were exported from BIORAD iQ5 software.

### Terminal restriction fragment (TRF) Southern blot assay

Animals were grown on 100 mm Petri dishes with NGM seeded with OP50. Synchronized adult animals were harvested and washed four times with M9 buffer. Pelleted animals were lysed for four hours at 50°C in buffer containing 0.1 M Tris-Cl (pH 8.5), 0.1 M NaCl, 50 mM EDTA (pH 8.0), 1% SDS, and 0.1 mg/mL proteinase K. DNA was isolated by phenol extraction and ethanol precipitation. DNA was eluted with buffer containing 10 mM Tris (pH 7.5) and 1 mM EDTA. DNA was then treated with 10 *μ*g/mL boiled RNase A. DNA was again isolated with phenol extraction and ethanol precipitation. Five *μ*g of DNA was digested with *Hinf*I at 37°C overnight. Telomere restriction fragment was blotted as described previously (Seo *et al.* 2015) (Supplementary Figure 14). Digoxigenin-labeled (TTAGGC)_4_ oligonucleotides were used as probes. Digoxigenin probes were detected with DIG nucleic acid detection kit (Roche). Blots were imaged with ImageQuant LAS4000 (GE healthcare).

### Fluorescence *in situ* hybridization assays

Fluorescence *in situ* hybridization (FISH) was performed as previously described (Seo *et al.* 2015). Embryos were isolated by bleaching synchronized adult animals using standard methods (Stiernagle 2006). Isolated embryos were fixed in 2% paraformaldehyde (PFA) for 15 minutes at room temperature (RT) on a polylysine treated glass slide. The slide was put on dry ice and freeze-cracked. The embryos were permeabilized in ice-cold methanol and acetone for 5 minutes each. The slides were washed with 1X phosphate buffered saline containing 0.1% Tween-20 (PBST) three times for 15 minutes each at room temperature. 10 *μ*l of hybridization buffer (50 nM Cy3-(TTAGGC)_3_ peptide nucleic acids probe (PANAGENE), 50% formamide, 0.45 M sodium chloride, 45 mM sodium citrate, 10% dextran sulfate, 50*μ*g/mL heparin, 100*μ*g/mL yeast tRNA, 100*μ*g/mL salmon sperm DNA) was added on the slide. The samples were denatured on a heat block at 85°C for three minutes. After overnight incubation at 37°C, the samples were washed in the following order: 1X PBST once for five minutes at room temperature, 2X SSC (0.3 M sodium chloride, 30 mM sodium citrate) in 50% formamide once for 30 minutes at 37°C, 1X PBST three times for 10 minutes each at room temperature. The samples were incubated in DAPI and mounted in anti-bleaching solution (Vectashield). The samples were imaged with a confocal microscope (LSM700, Zeiss). Telomere spots were quantified with TFL-TELO software (Dr. Peter Lansdorp, Terry Fox Laboratory, Vancouver) (Poon SS1, Martens UM, Ward RK 1999).

### Genome-wide association (GWA) mapping

GWA mapping was performed on marker genotype data and telomere-length estimates using the rrBLUP package (version 4.3) (Endelman 2011) and GWAS function. rrBLUP requires a kinship matrix and a SNV set to perform GWA. We generated a kinship matrix using our imputed SNV set with the A.mat function within rrBLUP. Genomic regions of interest were determined empirically from simulating a QTL that explained 20% of the phenotypic variance at each marker in our mapping data set. All simulated QTL were mapped within 100 markers (50 markers to the left and 50 markers to the right) of the simulated marker position. To generate a SNV set for mapping, we again used our imputed SNV set. However, we filtered the number of SNVs to a set of 38,688 markers. This set was generated by lifting over (from WS210 to WS245) a set of 41,888 SNVs previously used for GWA mapping (Andersen *et al.* 2012) and filtering our imputed SNVs to those sites.

### Million Mutation Project analysis

Whole-genome sequence data from mutagenized strains within the Million Mutation Project (MMP) was obtained from the sequence read archive (SRA, project accession number SRP018046). We removed 59 strains that were contaminated with other strains. We also were unable to locate the sequence data for 12 MMP strains on SRA, leaving us with a total of 1,936 mutagenized strains. Within the MMP project, read lengths varied among sequencing runs, being either 75 bp or 100 bp. We ran TelSeq on all sequencing runs assuming 100 bp reads. To utilize 75 bp sequencing runs, we took the 448 strains that were sequenced at both 75 and 100 bp lengths and used those estimates to develop a linear model. Then, this model was used to transform 75 bp length estimates to 100 bp estimates (Supplementary Figure 15). We then used the weighted average of telomere length estimates for all runs of a given strain based on total reads to calculate telomere-length estimates. Because telomeric reads resemble PCR duplicates, TelSeq utilizes them in calculating telomere length. However, we observed very low PCR and optical duplicate rates among MMP sequence data likely due to differences in library preparation in contrast to wild isolate sequence data. These differences likely account for shorter telomere estimates from the MMP sequence data.

Long-telomere strains from the MMP were classified as strains with telomere lengths greater than the 98^th^ quantile of all MMP strains (6.41 kb). Mutation data was obtained from the MMP website (http://genome.sfu.ca/mmp/mmp_mut_strains_data_Mar14.txt). A hypergeometric test was performed to identify which genes were enriched for mutations from long-telomere strains (Supplementary File 11) using the *phyper* function in R (R Core Team 2013). FX1400 was propagated for ten generations prior to whole-genome sequencing. Telomere length was estimated using TelSeq.

### Statistical analyses

Statistical analyses were performed using R (version 3.2.3). Plots were produced using ggplot2 (version 2.0.0).

### Longevity assays

At least 80 fourth larval stage animals were plated onto each of three separate 6 cm NGMA plates in two independent assays and viability assessed each day until all animals were scored as dead or censored from the analysis as a result of bagging or missing animals. Animals were scored as dead in the absence of touch response and pharyngeal pumping. Animals were transferred to fresh plates every day from the initiation of the assay until day seven of adulthood to remove progeny and transferred every other day until the completion of the assay. The following short telomere strains were scored: EG4349, JU2007, NIC1, and NIC3. The following long-telomere strains were scored: KR314, NIC207, QX1212, and RC301. Additionally, N2 and CB4856 were scored.

### High-throughput fecundity assays

Assays were performed similar to previously reported (Andersen *et al.* 2015) with the following differences. Animals were bleached, synchronized, and grown to L4 larvae in 96-well plates. From the L1 to L4 stage, animals were fed 5 mg/mL of a large-scale production HB101 lysate in K medium (Boyd *et al.* 2010) to provide a stereotyped and constant food source. Then, three L4 larvae from each of the 152 genotypes were dispensed using a COPAS BIOSORT instrument to wells containing 10 mg/mL HB101 lysate in K medium and progeny were counted 96 hours later. Fecundity data were calculated using 12 samples – triplicate technical replicates from four biological replicates. The data were processed using COPASutils (Shimko and Andersen 2014) and statistically analyzed using custom R scripts.

### Clustering of relatedness

Variant data for dendrogram comparisons were assembled by constructing a FASTA file with the genome-wide variant positions across all strains and subsetting by regions as described. MUSCLE (version v3.8.31) (Edgar 2004) was used to generate neighbor-joining trees. The R packages ape (version 3.4) (Paradis *et al.* 2004) and phyloseq (version 1.12.2) (McMurdie and Holmes 2013) were used for data processing and plotting.

## Acknowledgements

We would like to thank Joshua Bloom and members of the Andersen laboratory for critical comments on this manuscript. We also thank M. Barkoulas, T. Bélicard, D. Bourc’his, N. Callemeyn-Torre, S. Carvalho, J. Dumont, L. Frézal, C.-Y. Kao, L. Lokmane, I. Ly, K. Ly, A. Paaby, J. Riksen, G. Wang for isolating new wild *C. elegans* strains. The National Bioresource Project provided the FX1400 strain, and Wormbase data made a variety of analyses possible. This work was supported by an NIH R01 subcontract to E.C.A. (GM107227), the Chicago Biomedical Consortium with support from the Searle Funds at the Chicago Community Trust, and an American Cancer Society Research Scholar Grant to E.C.A. (127313-RSG-15-135-01-DD), along with support from the National Science Foundation Graduate Research Fellowship to D.E.C. (DGE-1324585).

